# Uumarrty and the Nash Score: a Game-theoretic, Agent-based Framework for Understanding Evolutionary Stability in Behavioral Traits

**DOI:** 10.1101/2023.08.09.552686

**Authors:** Michael Remington, Rulon W. Clark, Ryan J. Hanscom, Timothy E. Higham, Jeet Sukumaran

## Abstract

This paper introduces a new simulation framework for testing hypotheses relating to behavior strategies in predator-prey systems. To this end, we present two tools for simulating and analyzing behavioral trait dynamics: The Nash Score, a novel metric for evaluating evolutionary stability, and Uumarrty, an agent-based framework for simulating predator-prey interactions using game theory. These tools provide an approach for assessing the temporal co-evolution of behavioral traits within agent-based models, with a particular focus on predator-prey dynamics, though the framework is generalizable to other ecological interactions. The Nash Score functions as an analog to the Evolutionarily Stable Strategy (ESS) from classical game theory, offering a quantitative index to assess the relative stability and resilience of behavioral traits under selection. We demonstrate the utility of these tools through a case study on the microhabitat preferences of kangaroo rats and rattlesnakes. Specifically, we explore the emergence and stability of optimal strategies across scenarios with: (1) heterogeneous energy yields among microhabitats, (2) differential strike success rates by microhabitat, and (3) the presence of a specialist predator. Our results highlight how microhabitat specialist predators can drive other predators in the system to specialize due to outcompeting generalists at a given population frequency; leading to behavioral strategy stability in the system. Our case studies also show how behavioral trait dynamics can greatly vary depending on if you treat the trait as a pure strategy versus a mixed strategy. Collectively, this framework enhances our ability to explore ecological and evolutionary responses to environmental change, supporting more robust and comparable simulation-based research in eco-evolutionary dynamics.

## Introduction

When modeling predator-prey behavioral interactions, explicitly accounting for the co-evolution of both populations is important for accurately understanding of the system compared to modeling one of the populations as a fixed entity [1, 5, 15]. This phenomenon is due to the fitness landscape of both populations is interdependent on the behavioral strategy of the other species leading to a coupling effect [9]. However, modeling the co-evolution of both populations can be difficult from computational perspective as it introduces more variables and parameters, and increases the complexity of the output of the model as researchers need to summarize both populations. Our new method we propose in this paper that we term “Nash Score”, which is a post-hoc evolutionary stability analysis we use to summarize the temporal co-evolutionary effects of behavioral trait evolution in an agent-based system.

The Nash Score functions as an analog to the concept of the evolutionarily stable strategy (ESS) in evolutionary game theory [24], but for agent-based modeling framework. This new method will give researchers a standardized metric to assess the evolutionary stability of traits in an arbitrarily complex, co-evolving, simulated systems under various conditions. The Nash Score also can be utilized as a measure of the degree of divergence from the stable point, as it is a continuous metric. This provides a standardizes, quantifiable description of the system under a given set of conditions. From a theoretical perspective, the Nash Score can be used to tell whether the system is either periodic or bi-stable which is typical of evolutionary arms races, versus the system state reaching the ESS.

Taking tools from evolutionary game theory and integrating them into an agent-based system addresses several different gaps in the field of modeling predator-prey interactions. Firstly, utilizing agent-based models addresses critiques of game theory being too simplistic and unable to capture the complexity of real world systems, especially across biological scales [16]. Secondly, it provides a standardized post-hoc analysis of agent-based systems which will allow for the ability to compare and contrast the results of simulation studies and non-linear systems among researchers which has been a critique of agent-based modeling [3, 8, 12]. Finally, it allows researchers to better understand the emergent consequences of changes in ecological patterns, to eco-evolutionary patterns which is useful for better understanding the long term consequences of changing environmental conditions [11].

To demonstrate how our methods can contribute to a better understanding of the evolution of optimal behavioral strategies, we apply our approach to a predator-prey system of a generalist prey (kangaroo rats) and a generalist sit-and-wait predator (rattlesnakes) to assess how they influence one another’s microhabitat preferences under various experimental designs. We use a game theoretic agent-based model we entitled “Uumarrty” along with the Nash Score to re-create a previous mathematical game theory model which analyzed the same system to see the effects of a secondary predator (owls) on microhabitat usage behavioral strategies of kangaroo rats and snakes [5]. They found the conditions that lead to an ESS was when the risk from owls in the open is larger than the risk from snakes in the bush, all snakes and all rodents should select the bush microhabitat. Thus if our model is valid, by adding a specialist predator (owls) into our system, we expect to observe similar qualitative findings of the optimal behavior strategy and whether strategy reaches an ESS.

In conjunction with our case study of introducing a specialist predator into the system, we explore two other experimental designs; first, we assess the influence of increasing heterogeneity between microhabitat energy gains from foraging for the kangaroo rats. This case study could be viewed as a bottom-up effect the selection pressure only directly impacts the prey. Secondly, we analyze how increasing heterogeneity between microhabitat dependent strike success probability for snakes. This could be seen as a top down effect as we assume strike success is intrinsic to the generalist predator.

A long-standing debate revolves around whether the microhabitat preferences of prey are primarily driven by risk aversion caused by predator decisions [7], or by decisions made by prey to maximize foraging yields [25]. One primary objective of all of our experimental designs is to determine the ecological factors that most influence shifts and stability in the microhabitat preferences of a predator-prey system. By comparing the Nash Score analysis results across this set of case studies, we can come of what are important factors driving behavior of organism’s of the system as a whole.

The dynamics of microhabitat usage in rattlesnake and kangaroo rat systems have been studied in both empirical [4, 6, 10, 14, 22] as well as theoretical contexts such as the game theory model described earlier [5], an agent-based model [21], and other mathematical frameworks [7]. This makes this system suitable for comparing our methodology to a rich body of previous research. The goal of this paper is to confirm the validity of the use of the Nash Score, as well as to showcase how researchers can utilize the Nash Score for novel analysis of community dynamics as a whole.

## 1 Methods

### 1.1 Construction of the Model

#### 1.1.1 Agent-based Modeling and the Genetic Algorithm

Our model, which we termed “Uumarrty”, is an agent-based, stochastic predator prey model written in Python 3.8 [26]. The basic model has two main populations of agents: the kangaroo rat agent (prey) and the rattlesnake agent (predator). For the populations of agents, we applied an evolutionary genetic algorithm to both populations that acts on a microhabitat preference trait that is assigned to individuals of the population at birth (Figure 1). Agents move in a virtual landscape composed of individual cells representing microhabitats based on a random decision that is weighted by the behavioral preference trait for one particular type of microhabitat over another. This microhabitat preference trait is an inherited, mutable trait assigned at birth of the new agent. These virtual organisms are modeled as asexual populations with discrete, non-overlapping generations.

**Fig 1.**
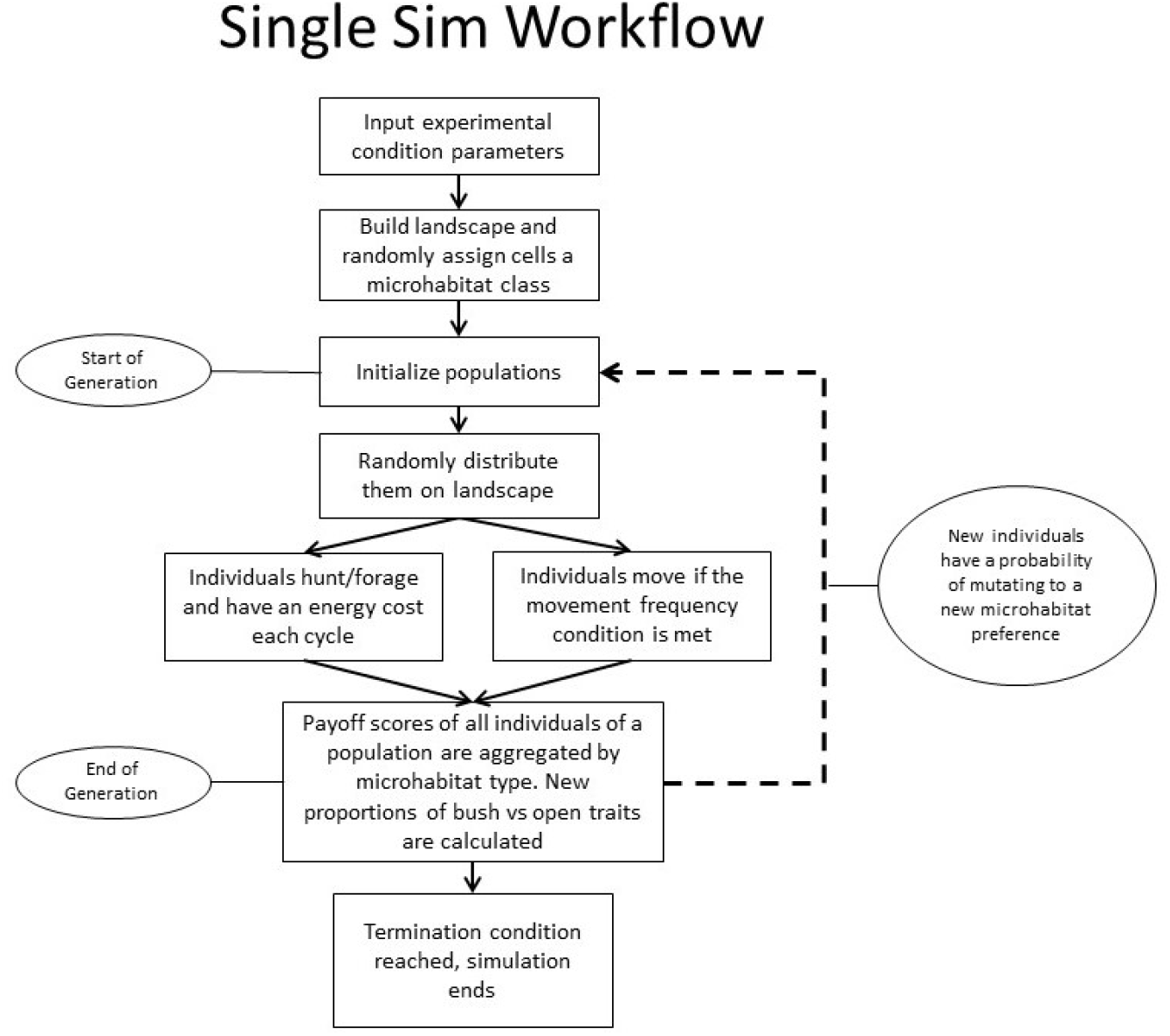
Conceptual diagram showcasing the logic of an individual simulation. This logic is applied to both the kangaroo rat population and the rattlesnake population of all simulations.

During an agent’s generation time, agents accrue payoffs by successfully foraging without dying from either predation or energy depletion. Organisms are able to move within a generation but their preference is treated as a constant trait that is stationary within generation, but evolves over the course of the simulation. There are 3 parameters set by the user (owl movement frequency, snake movement frequency, and kangaroo rat movement frequency) which indicates how many cycles pass before they move to a new location. See Table 1 for parameter values. Individuals that accrue a higher pay-off compared to others in the population are able to leave more offspring with the same microhabitat preference unless a random mutation occurs. At the end of each generation time, the pay-off scores from the set of agents who survived are aggregated by their microhabitat preference type. The proportion of the sum of pay-off scores of the individuals with a given microhabitat preference trait over the total sum of pay-off scores of all agents in the species class is then the proportion of agents in the next generation that will retain the same trait. Each population persists on the landscape for a set number of feeding and/or movement cycles which is also referred to in the model as a generation (Figure 1).

**Table 1.** Constant experimental parameters. Cells titled “EXP” are parameters that are associated with experimental groups for a given case study. Parameters that state “per x cycles” indicate that the action is done once after x number of cycles in the associated cell have passed.

| Parameter | Owl<br>Exp | Energy<br>Gain<br>Exp | Snake<br>Strike Exp |
| --- | --- | --- | --- |
| Cycles | 500,000 | 500,000 | 500,000 |
| Landscape Size X Direction (cells) | 25 | 25 | 25 |
| Landscape Size Y Direction (cells) | 25 | 25 | 25 |
| Proportion Open Habitat | 50% | 50% | 50% |
| Proportion Bush Habitat | 50% | 50% | 50% |
| Kangaroo Rat Population Size | 500 | 500 | 500 |
| Rattlesnake Population Size | EXP | 100 | 100 |
| Owl Population Size | EXP | 0 | 0 |
| Per-cycle Snake Death Probability | 0.001 | 0.001 | 0.001 |
| Snake Strike Success Probability<br>(Bush) | 0.21 | 0.21 | EXP |
| Snake Strike Success Probability<br>(Open) | 0.21 | 0.21 | EXP |
| Owl Strike Success Probability (Bush) | 0.0 | NA | NA |
| Owl Strike Success Probability (Open) | 0.21 | NA | NA |
| Kangaroo Rat Energy Gain (Bush) | 12 | EXP | 12 |
| Kangaroo Rat Energy Gain (Open) | 12 | EXP | 12 |
| Snake Energy Gain | 1500 | 1500 | 1500 |
| Kangaroo Rat Energy Cost | 7 | 7 | 7 |
| Snake Energy Cost | 14 | 14 | 14 |
| Kangaroo Rat Movement Frequency<br>(per x cycles) | 1 | 1 | 1 |
| Snake Movement Frequency (per x<br>cycles) | 8 | 8 | 8 |
| Owl Movement Frequency (per x<br>cycles) | 1 | NA | NA |
| Kangaroo Rat Mutation Probability | 0.01 | 0.01 | 0.01 |
| Snake Mutation Probability | 0.01 | 0.01 | 0.01 |
| Kangaroo Rat Reproduction Frequency<br>(per x cycles) | 50 | 50 | 50 |
| Snake Reproduction Frequency (per x<br>cycles) | 300 | 300 | 300 |

The pay-off score is a proxy for fitness that is parameterized by the user of the software. In our model, the only populations that are capable of reproducing are the kangaroo rats and the rattlesnakes. However, when the trait is actually assigned to the individual of the next generation, there is some probability that the trait will mutate into another microhabitat preference strategy based on a set probability by the user. The kangaroo rat and rattlesnake populations interact for a set number of feeding cycles set by the user before the simulation is terminated and the results are aggregated into a format to be summarized (Figure 2).

**Fig 2.**
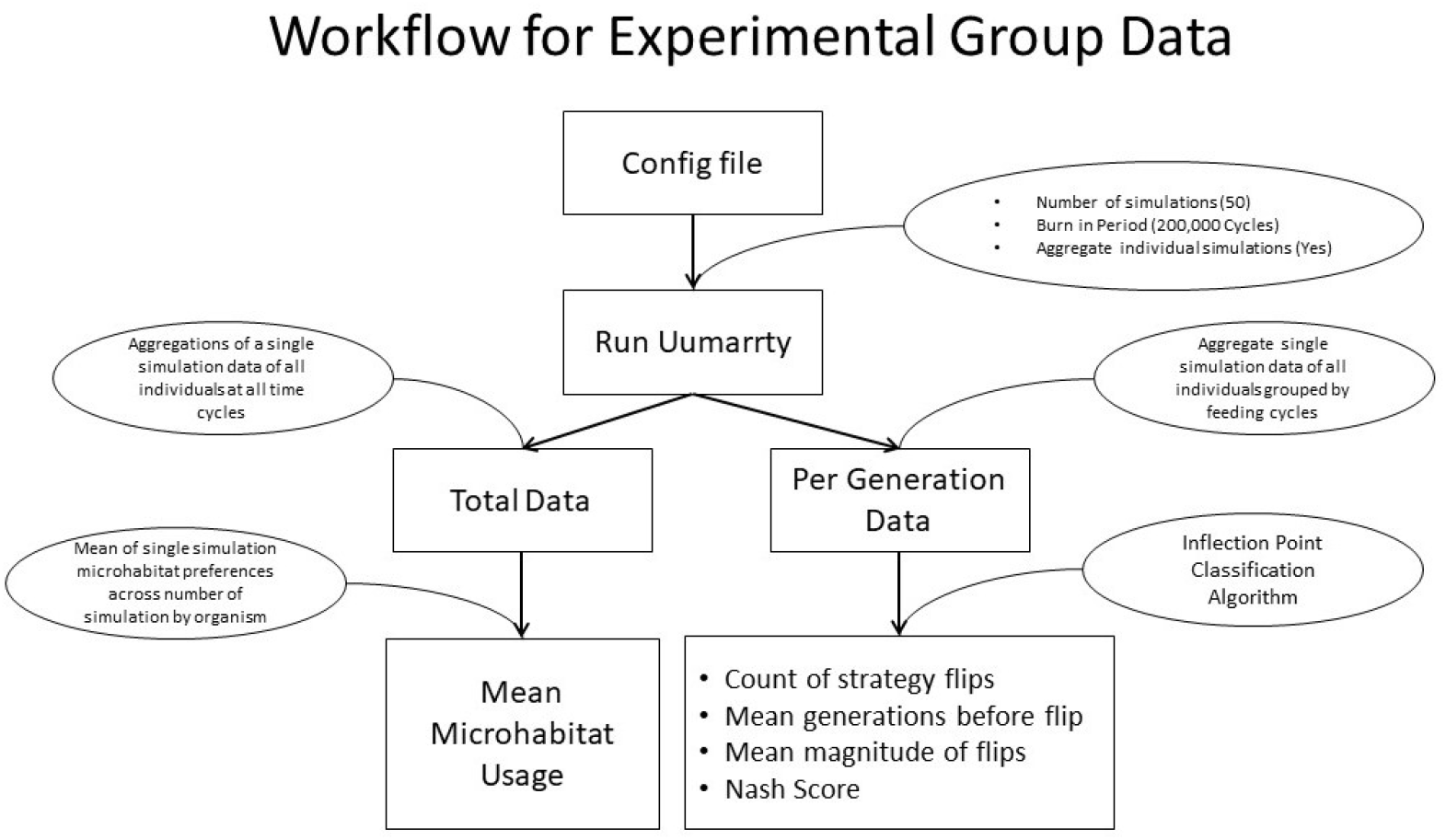
Conceptual diagram of the simulation workflow for how to generate the results of one experimental group of a case study. This work flow was applied to all 6 experimental groups of all 3 case studies.

Population sizes are semi constant. At the start of each generation, populations with the genetic algorithm applied to them start at an initial population size parameterized by the user. Over the generation time, individuals are removed from the simulation if they are successfully predated or starve. The user may also set a variable for a random chance of dying to model the other ecological processes that may affect this population outside the scope of the simulation such as predation or disease. For example, we set a parameter in our case study called “Per-cycle Snake Death Probability” which is a 0.1% chance per cycle that the snake dies (Table 1). At the end of the generation, pay off scores are aggregated among the surviving individuals and the next generation is reset back to the initial population size with the population’s proportions of traits being dependent on the success of the individuals of the previous generation.

#### 1.1.2 Landscape

Kangaroo rats and rattlesnakes predominately reside in sparsely vegetated arid environments with significant open space. To simplify this environment in silico, we modelled the landscape as a two dimensional grid composed of two classes of cells that represent microhabitats; the bush and the open. The bush microhabitat represents any habitat that gives an aerial canopy cover such as creosote (*Larrea tridentata*) or mesquite (*Prospis spp*.). The open microhabitat represents areas with sparse vegetation such as grasses and with no aerial cover. All microhabitat cells of the same class are equal in terms of parameterization and agents operate and interact in these cells. The interactions in a given microhabitat vary based on the classification of the cell and the input parameters of the model. Pay-offs for foraging in these microhabitats are then accrued by a prey individual based on the microhabitat they occupy during a feeding event. The specific parameters we use in our model can be seen in Table 1.

Experimental parameters such as the number of owls (specialized aerial predators that only forage in the open), energy gain from prey foraging in a given microhabitat, and the probability of a predator’s strike being successful in a microhabitat are set as initial conditions of the model (Table 2, 3, 4).

**Table 2.** Experimental parameters for the case study of differing foraging energy gains in microhabitats for Kangaroo Rats.

| Experiment | Energy Gain (Open) | Energy Gain (Bush) |
| --- | --- | --- |
| Group 1 | 12 | 12 |
| Group 2 | 14 | 10 |
| Group 3 | 16 | 8 |
| Group 4 | 18 | 6 |
| Group 5 | 20 | 4 |
| Group 6 | 22 | 2 |

**Table 3.** Experimental parameters for the third case study of differing strike success experimental groups.

| Experiment | Strike Success Probability (Bush) | Strike Success Probability (Open) |
| --- | --- | --- |
| Group 1 | 0.21 | 0.21 |
| Group 2 | 0.231 | 0.189 |
| Group 3 | 0.252 | 0.168 |
| Group 4 | 0.273 | 0.147 |
| Group 5 | 0.294 | 0.126 |
| Group 6 | 0.315 | 0.105 |

**Table 4.**
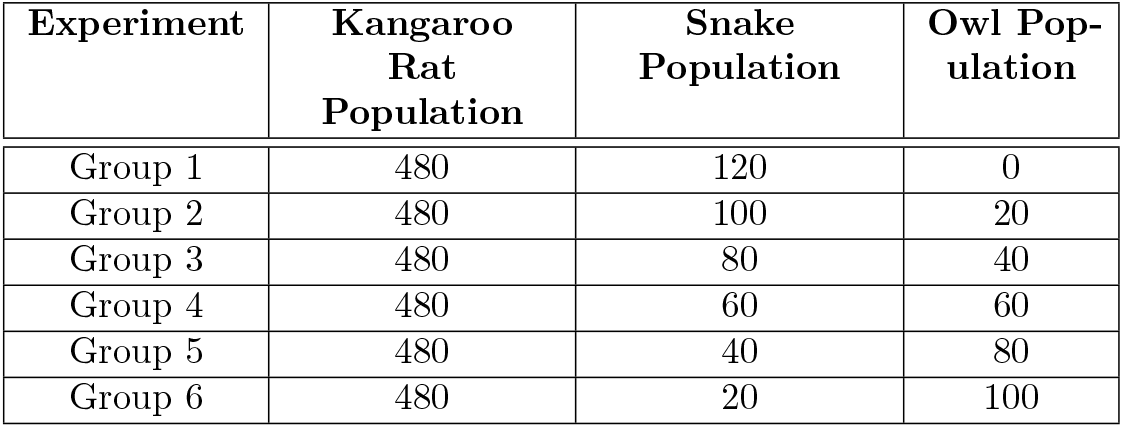
Experimental parameters for the case study of introducing owls into the system experimental groups.

| Experiment | Kangaroo Rat Population | Snake Population | Owl Population |
| --- | --- | --- | --- |
| Group 1 | 480 | 120 | 0 |
| Group 2 | 480 | 100 | 20 |
| Group 3 | 480 | 80 | 40 |
| Group 4 | 480 | 60 | 60 |
| Group 5 | 480 | 40 | 80 |
| Group 6 | 480 | 20 | 100 |

#### 1.1.3 Classes of Agents

There are two classes of agents for our model, the predator class and the prey class. For both classes, the goal of the agent is to maximize their pay-off score and avoid dying. The pay-off score for each class of agent is a running score that accrues throughout it’s lifetime per cycle. Equation 1 represents how the energy score is calculated each cycle for each individual. In Equation 1, *p*_*n*_ is the current energy score of the individual, *α* represents the energy cost per cycle, and *ω* represents the energy gain. For both classes of agents, *α* is a constant. Energy gain *ω*, differs between the two classes of agents and for prey agents, can also vary by microhabitat.

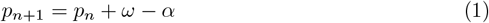

For predators, *ω* is dependent on an interaction with a prey agent and a strike success probability set as a parameter by the user that can differ based on microhabitat. If a strike is successful, the prey agent is removed from the landscape and a energy amount *ω* accrues to the predator agent’s pay-off score. For prey agents, *ω* is a constant that is dependent on the type of microhabitat the organism occupies. For both classes of agents, at the beginning of each cycle, if *p*_*n*_ is less than 0, then the agent is considered to have starved and is removed from the environment. In our simulations, pay-off values for snake energy gain, snake energy cost, kangaroo rat energy gain (bush/open), and kangaroo rat energy cost were based on a previous game theory model analyzing the same system [5].

#### 1.1.4 Pure and Mixed Strategies

Uumarrty is capable of modeling the microhabitat preference of an individual as a pure strategy or a mixed strategy [24]. A pure strategy is where an individual exclusively prefers one microhabitat over another, while a mixed strategy is where individuals have a weighted preference for one microhabitat over another, but can still utilize both microhabitats for foraging over the course of their lifetime.

Equation 2 represent the probability of an individual moving into a given cell in a landscape. Microhabitat preference (*mp*) is the individual’s weighted preference of a given microhabitat, *bc* is the total number of bush microhabitats in the landscape, *oc* is the number of open cells in the landscape, *i* is an individual bush cell in the landscape, and *j* is an individual open microhabitat cell. Both the pure and the mixed strategies use these sets of equations for movement, however; the parameter *mp* is binary when microhabitat preference is modeled as a pure strategy, and continuous from 0 to 1 (rounding to two decimal places) when microhabitat preference is modeled as a mixed strategy. The *mp* value of 1 represents a complete preference for the bush and a *mp* value of 0 represents a preference for the open. This value is passed down to the offspring with a probability of mutating. If a mutation event occurs, the magnitude of the mutation is sampled from a Gaussian distribution with a standard deviation set as a parameter of the model.

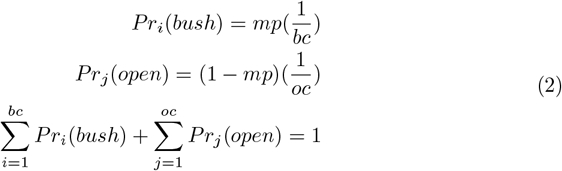

### 1.2 The Nash Score and Post Simulation Analysis

After 50 simulations for each experimental group, our model summarizes data for both the total run and per generation (Figure 2). The data for the total run is a global mean of all the microhabitat preference traits of all individuals of a population at every time cycle. These means are compared across experimental groups to evaluate strategy choices. This value is the optimal strategy of the population under the set experimental conditions. A one way ANOVA and Tuckey post hoc test for multiple comparisons are implemented to establish statistically significant differences between experimental groups of a case study. Statistical tests were conducted using the statsmodels package in Python [23].

The simulation software also summarizes mean microhabitat preference of a population aggregated by each feeding cycle. To convert this to per generation, only the last cycle of the generation time is used and the other data mid generation is filtered out. Under certain selection conditions for a given simulation, microhabitat preference per generation will fluctuate in a stochastic, oscillating manner (Figure 3A), or may become constant (Figure 3B). We developed a novel method to classify the local optima points of the per generation microhabitat preference strategies in order to obtain metrics for stability analysis.

**Fig 3.**
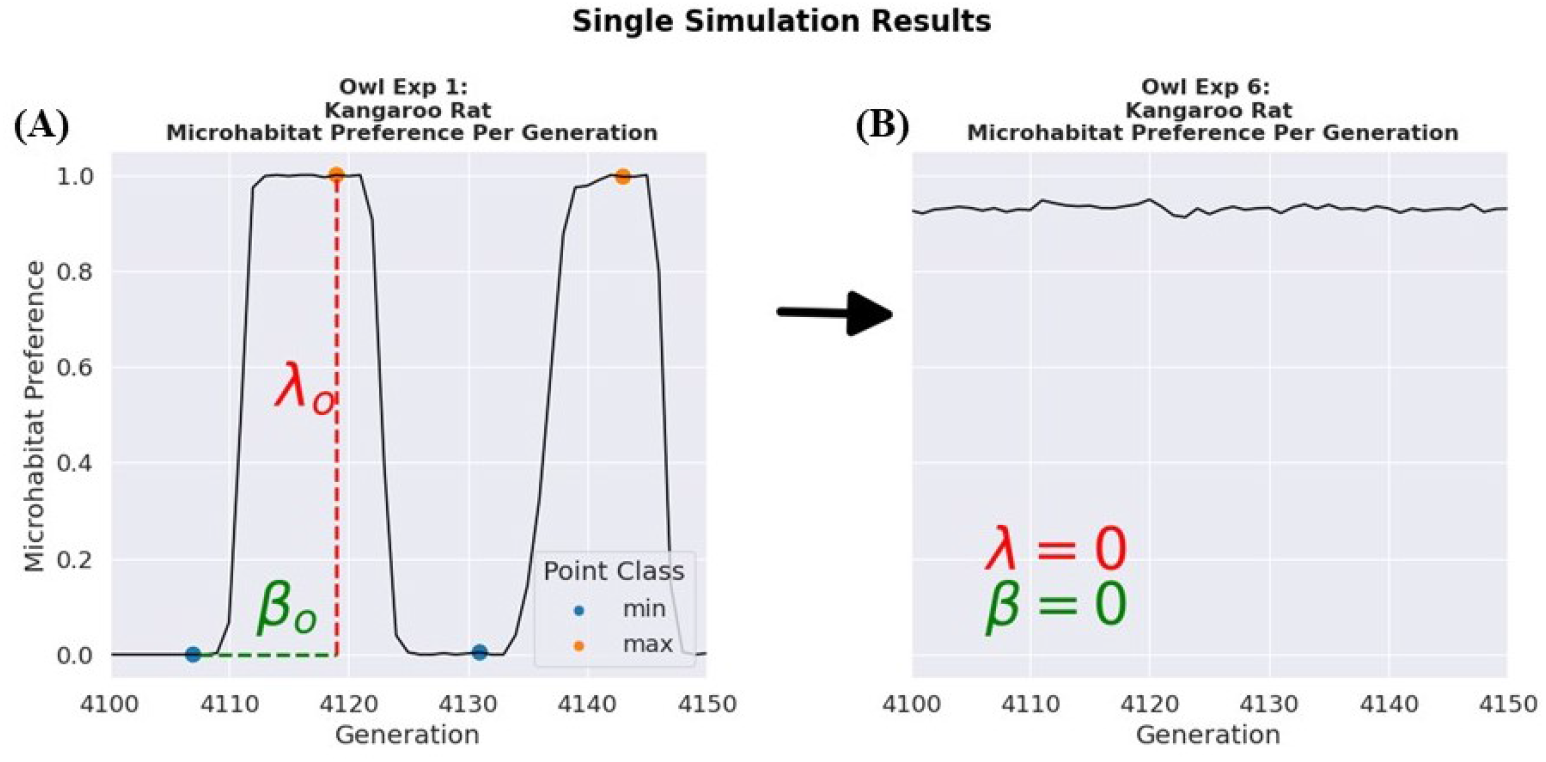
Both graphs are the mean microhabitat preference for the kangaroo rat population per generation during an individual simulation. A microhabitat preference of 1 is a complete preference toward bush microhabitats while a microhabitat preference of 0 is a complete preference toward the open. (A) is a single simulation for the owl experiment from experimental group one. (B) is a single simulation for experimental group 6. The metric *λ*_0_ is the difference between the microhabitat preference of an optimal point and the previous optimal point in the time series which represents the magnitude of the strategy flip, shown by the red dotted line. The metric *β*_0_ is the number of generations from one optimal point to the previous one. When *λ* and *β* are zero this means there are no optimality points and the microhabitat preference of the population is in equilibrium which can be seen in (B). Point class represents whether an optima point is a maximum or a minimum point. The arrow indicates how from experimental group 1 to experimental group 6, the microhabitat preference of the kangaroo rat population reached an equilibrium due to the introduction of owls.

Our local optima point classification algorithm works by using a central approximation of the first derivative, and the second derivative of the per generation data with a window size of three generations. Because both vectors are highly chaotic, we applied a Savitzky Golay smoothing function [20] with a window size of three. We used two methods to classify local optima points; the first is a traditional calculus method where minimums are defined when the first derivative of a point is equal to zero and the second derivative is positive. Maximums are classified if the second derivative is negative. However, this can be unreliable due to portions of the data that rapidly oscillate between positive and negative values. If the first method does not classify local optima point at a given time step, the algorithm checks if the first derivative changes from being negative at the current time step to being positive at the next time step with a magnitude greater than a set sensitivity, then this point would be classified as a minimum local optima point and the opposite would be a maximum. A sensitivity parameter of the absolute value of the second derivative being greater than 0.1 was used in the classification algorithm in order to only classify one local maxima point per strategy flip as rapid small changes of the data could produce a false local optima.

From a biological context, local optima points would represent populations undergoing a shift toward a different microhabitat preference strategy, likely as a result of interspecific competition or predation. From the local optima point classification algorithm, three descriptive statistics are generated to understand the dynamics of strategy flip oscillations: the total count of strategy flips, the mean number of generations before a strategy flip, and the mean magnitude of a strategy flip (Figure 3A). Unstable microhabitat preferences are characterized by having a large number of strategy flips, a low value for mean generations before a strategy flip, and a larger value for the mean magnitude of strategy flips. We classify a completely evolutionary stable strategy for microhabitat preference as having zero local optima points and being a constant trend. However, general stochasticity from the model can lead to occasional false positive local optima points. To account for this, the algorithm has sensitivity conditions. If there are less than five local optima points, and a mean magnitude before strategy flip value of less than 0.05, then the results of the per generation data of a given simulation is classified as stable 1 or unstable 0.

Due to the stochastic nature of an agent-based framework, there can be variance in the results of simulations run using the same set of experimental parameters. To quantify the stability of the microhabitat preference strategies as a whole for a given experimental design, we used the Nash Score. The Nash Score is the ratio of simulations classified as stable over the total number of simulations of an experimental group. A Nash Score of 1 would be analogous to the evolutionarily stable strategy used in game theory, as it demonstrates a consistent outcome of in all simulations at all generations of an experimental group, and mutant behavioral strategies cannot invade the fixed microhabitat preference adopted by a population. The advantage of the Nash Score is that it is a continuous metric, and can be used to assess and compare relative “evolutionary stability” across different simulation conditions. Stability is not likely to be a property of most evolutionary systems in nature, as they are subject to stochasticity from changing abiotic and biotic conditions, and a metric for evolutionary stability that is continuous is arguably more useful for cross-system comparisons than a categorical metric. A Nash Score of less than one but greater than zero would indicate the system is approaching an ESS and is semi-stable under a given set of experimental conditions.

From a theoretical perspective, this algorithm matches the definition of the ESS because under the genetic algorithm, new mutants will always arise in the population. In cases where behavioral trait strategies are constant and no inflection points can be identified, all mutants that deviate from the optimal strategy have such comparatively low fitness or survivability that their microhabitat preference has no effect on the populations mean preference due to strong selection against mutant traits. A Nash Score of 1 would indicate the fact that no mutant strategies can invade the optimal behavioral preference strategy, thus it can be classified as an ESS. A Nash Score of less than one but greater than zero would indicate the system is approaching the ESS. This gives the user of the model a sense of how certain selection pressures lead to behavioral traits becoming evolutionarily stable strategies.

### 1.3 Kangaroo Rats vs. Rattlesnakes

For each experimental group of all case studies described below, we modelled the interaction between kangaroo rat and rattlesnake populations with the goal of first identifying optimal microhabitat usage strategies in a basic predator-prey scenario then introducing more complexity into the system in subsequent experimental groups that follow. Each set of experiments is also run with a control group with consistent experimental parameters across all the case studies. In the experimental groups, model parameters are systematically altered and outcomes are compared to the control group to evaluate the evolution of microhabitat preferences of the population and how it may diverge from the expected frequency seen in the control group.

Input parameters for kangaroo rat and rattlesnake population agents were derived from ecological data. Rattlesnakes live about 5 times longer than kangaroo rats, so in our model the reproduction frequency of kangaroo rats after 50 foraging events while rattlesnakes reproduce after 300 foraging events. Rattlesnake strike success probability (1 in 4 encounters result in death of kangaroo rat) are from empirical data in [27]. Energy gain parameters are from a game theory model [5] that was in turn based on empirical field observations and experiments [4]. Parameters used in case studies are listed in Table 1.

#### 1.3.1 Case Study 1: Differing Foraging Energy Gains for Kangaroo Rats in Microhabitats

Different microhabitats may provide more nutritious or abundant food resources to the kangaroo rats, which is likely to influence microhabitat preference. To see the effect of how food resource levels may influence the microhabitat preference of the prey and it’s predator, we increase the energy gain for the kangaroo rats in one microhabitat over the other (Table 2). This experimental design should help researchers ascertain how the ecological impact of a bottom up regulation of energy gain from prey foraging influences the microhabitat preference of both the predator and prey.

#### 1.3.2 Case Study 2: Differing Strike Success of Rattlesnakes in Bush Microhabitats

Although empirical data is lacking, it is possible that the increased structural complexity of vegetated microhabitats helps conceal ambush-hunting rattlesnakes, thus leading to increased strike success toward prey. To simulate this scenario, we increased the strike success of rattlesnakes in the bush by 1%, while decreasing the strike success in the open by 1%, thereby holding total selection pressure constant, but varying risk across microhabitats (Table 3).

#### 1.3.3 Case Study 3: Introducing Owls

There have been several studies on the effects of owls on kangaroo rat and small mammal foraging [4, 5, 10] that generally show owl predation results in kangaroo rats decreasing foraging time in open habitats, where they are susceptible to aerial attacks. In this experiment, we introduce a risk of predations from owls, specialist predators that only utilize the open microhabitat for foraging. We assume they are specialist predators in the context of microhabitat usage due to the canopy cover providing protection to kangaroo rats in bush microhabitats. We also assume that owls are unable to predate rattlesnakes. The genetic algorithm applied to the kangaroo rats and rattlesnakes is not applied to the owl population. We hold the total number of predators constant but manipulate the ratio of snakes to owls in a landscape to see how this will influence the kangaroo rat and rattlesnake microhabitat preference (Table 4).

## 2 Results

For each experimental group, we obtain results from an ANOVA test and a Tukey-post hoc test (Figure 4) for both the rattlesnake and kangaroo rat population and; for pure and mixed microhabitat preference. The model output also summarizes mean microhabitat preference (Figure 5-6), the total amount of microhabitat usage strategy changes (Figure 7-8A,D,G), mean generations before a strategy flip (Figure 7-8B,E,H), and the mean magnitude of how severe the strategy change is (Figure 7-8C,F,I). Finally, we visualize Nash Score results as a heat map in Figure 9. For all the case studies, kangaroo rat and rattlesnake populations, the ANOVA results showcased a significant p-value showcasing that all of our experimental designs had significant effects on changing the microhabitat preferences of both populations compared to the control group. However, the results of the Tukey-post hoc test indicates that some experimental groups were more similar than others (Figure 4).

**Fig 4.**
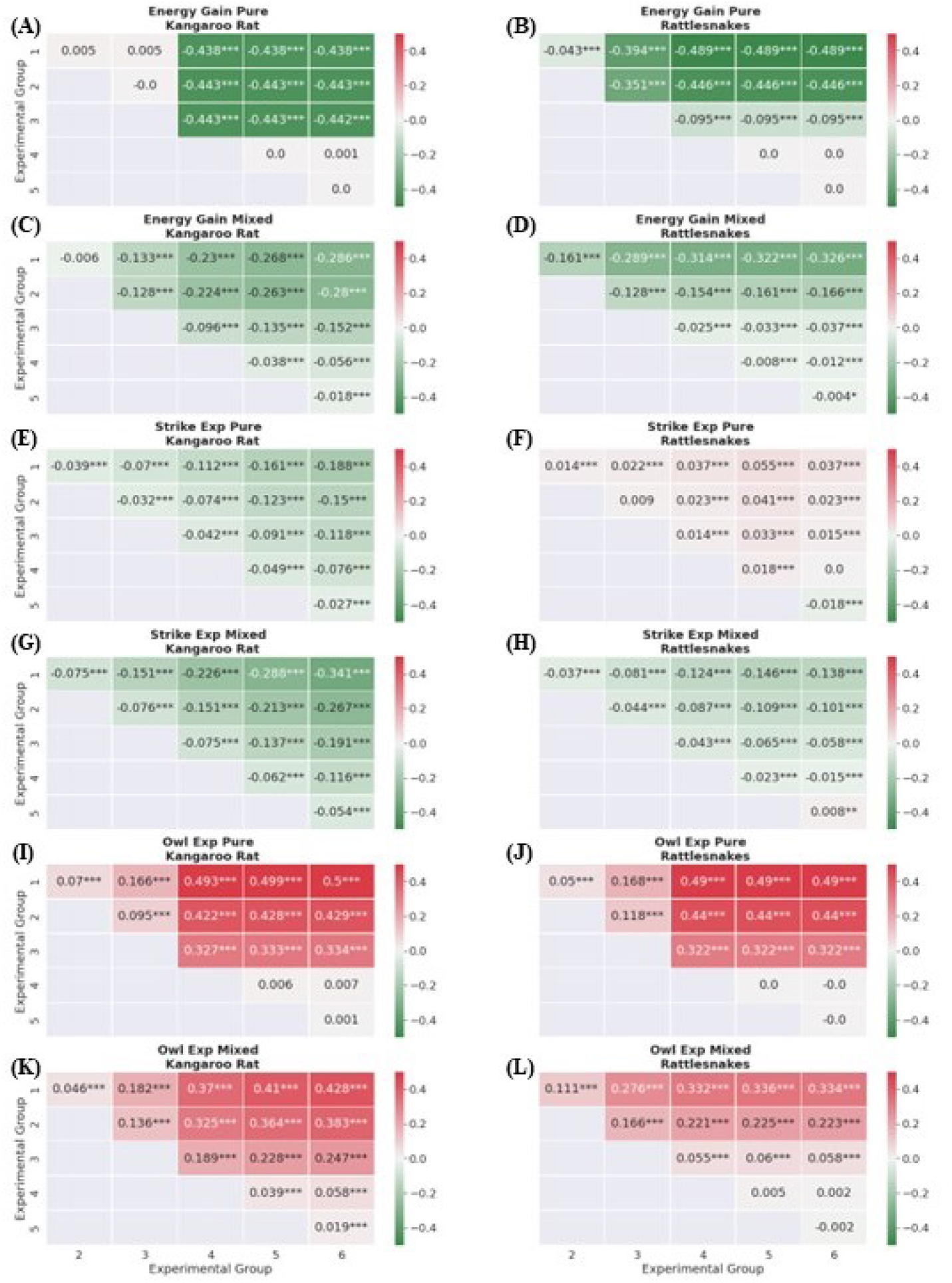
Results of the Tukey Post Hoc anova test. Values in each cell are the differences of the means between experimental groups. A positive increase (red) demonstrates an increased preference to the bush, while a negative mean difference indicates an increased preference for the open. Stars represent whether the experimental groups have statistically significantly different means. *** represents a p-value less than 0.001, ** represents a p-value less than 0.01, finally a p-value less than 0.05 is represented by a single *.

**Fig 5.**
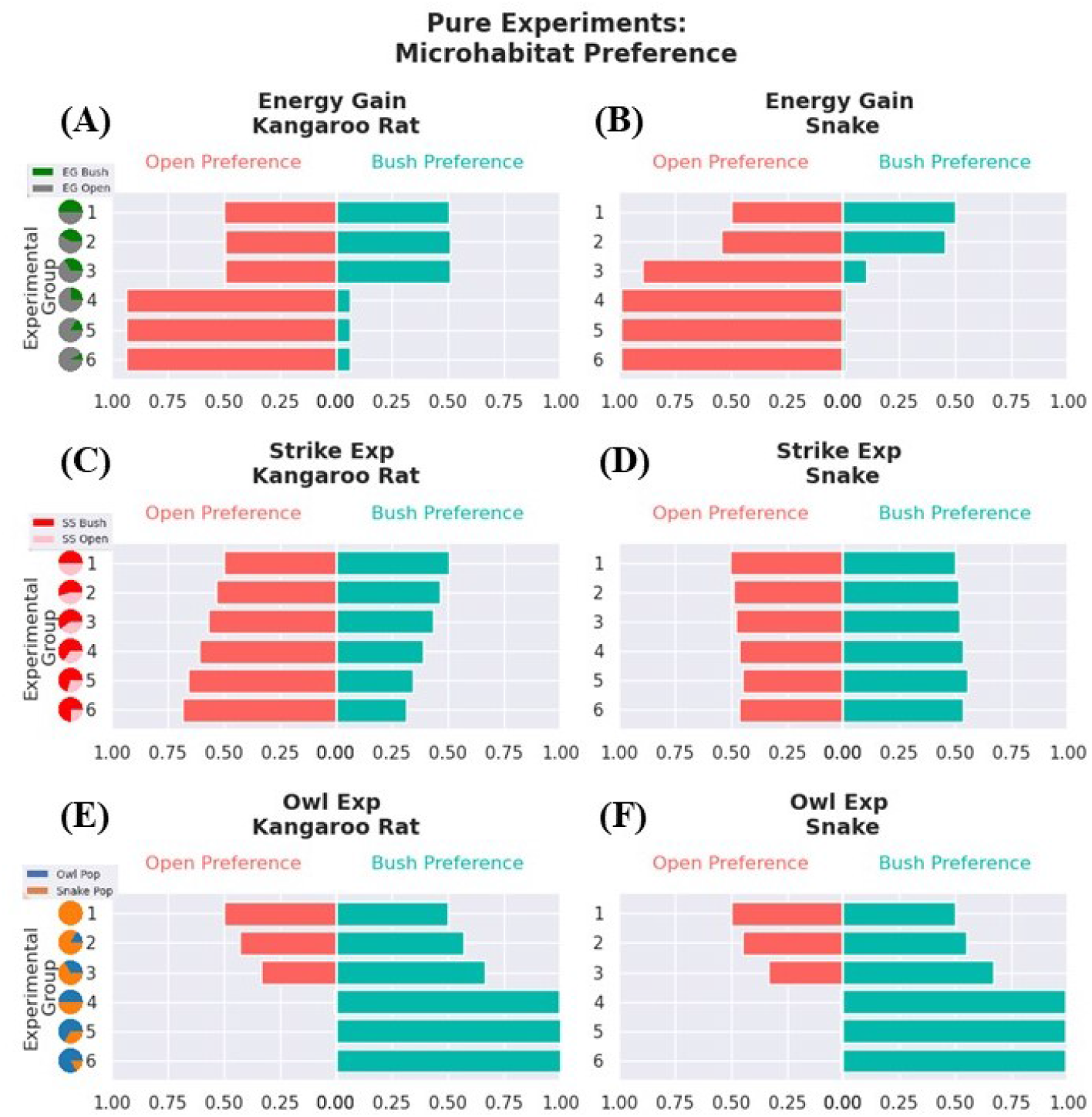
Mean microhabitat preference of the populations for all experiments with pure microhabitat usage strategies. The y-axis shows the experimental groups with pie charts representing how the experimental parameter of interest is being changed. The x-axis is the mean microhabitat preference of the population over all generations sampled from the simulation. The sum of the open preference bar graph (red) and the bush preference bar graph (blue) in all figures is equal to 1. (A) shows the microhabitat preferences of kangaroo rats for the differing foraging gains experiment for the kangaroo rat population. (B) is the same as (A) but for the rattlesnake population. Pie charts are of the values in Table 2. (C) is the microhabitat preferences of kangaroo rats in the rattlesnake strike probability experiment. Pie charts are of the values in Table 3. (D) is the same as (C) but for the rattlesnake population. (E) is the microhabitat preferences of the kangaroo rat population under different experimental designs for the owl experiment. Pie charts are of the values in Table 4. (F) is the same as (E) but for the snake population.

**Fig 6.**
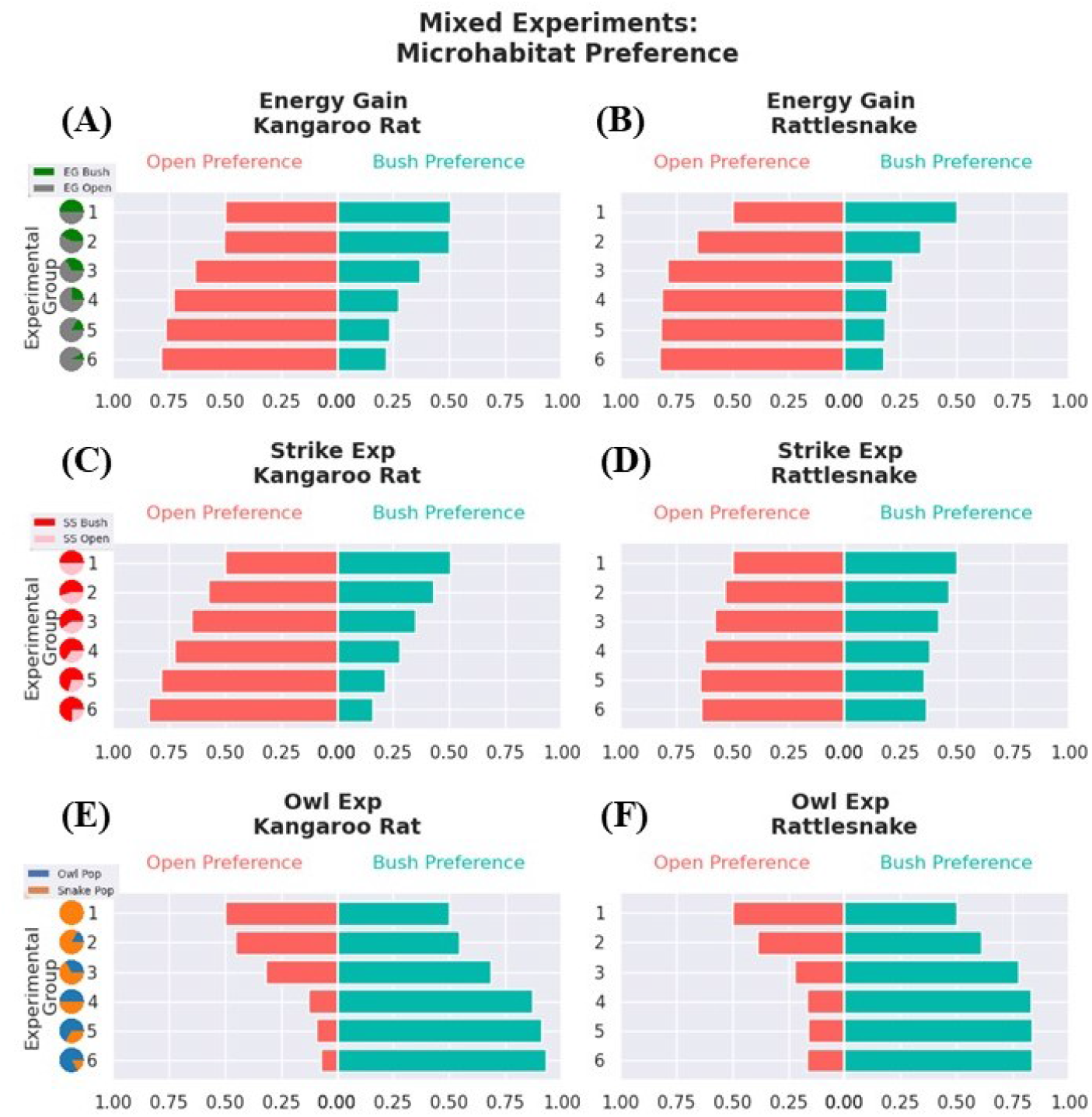
Mean microhabitat preference of the populations for all experiments with mixed microhabitat usage strategies. The y-axis shows the experimental groups with pie charts representing how the experimental parameter of interest is being changed. The x-axis is the mean microhabitat preference of the population over all generations sampled from the simulation. The sum of the open preference bar graph (red) and the bush preference bar graph (blue) in all figures is equal to 1. (A) shows the microhabitat preferences of kangaroo rats for the differing foraging gains for kangaroo rats in the two microhabitats. Pie charts are from the values in Table 2. (B) is the same as (A) but for the rattlesnake population. (C) The microhabitat preferences of kangaroo rats in the rattlesnake strike probability experiment. Pie charts are from the values in Table 3. (D) is the same as (C) but for the rattlesnake population. (E) is the microhabitat preferences of the kangaroo rat population under different experimental designs for the owl experiment. Pie charts are of the values in Table 4. (F) is the same as (E) but for the snake population.

**Fig 7.**
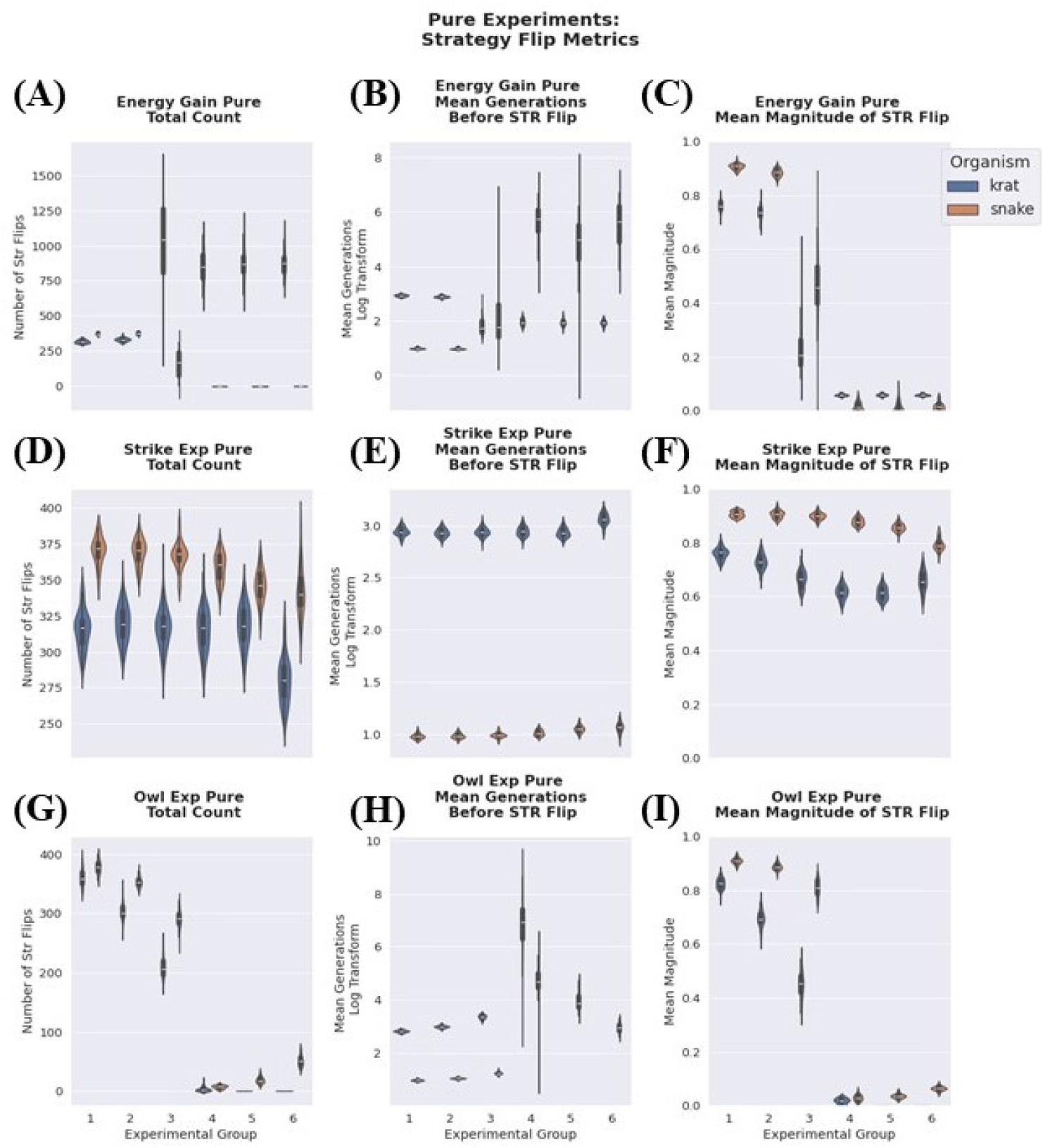
Analysis of metrics produced by the point classification algorithm such as total number of strategy flips, the mean number of generations before a strategy flip, and the mean magnitude of a strategy flip. All of these figures are for the pure microhabitat usage trait.

**Fig 8.**
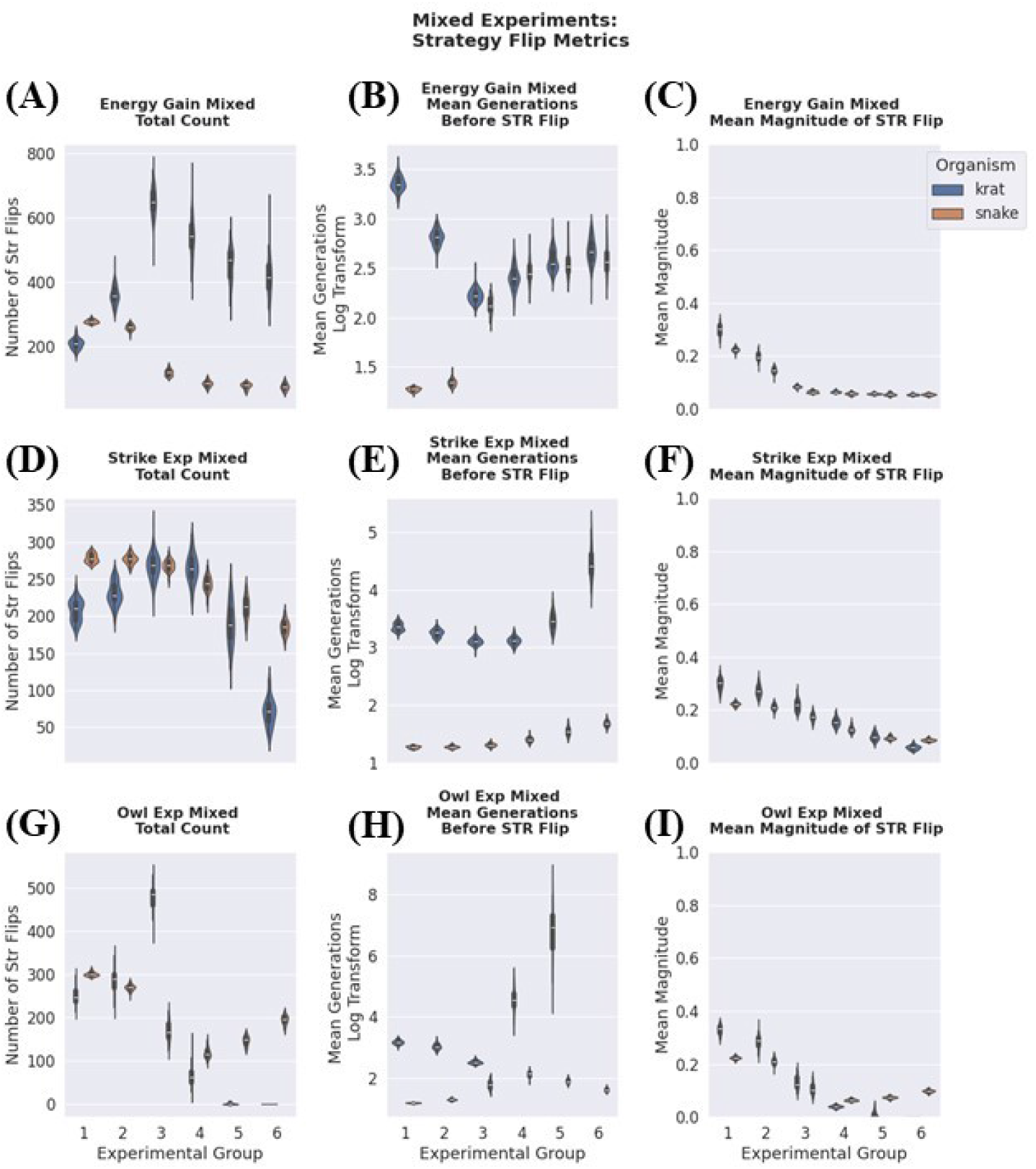
Analysis of metrics produced by the point classification algorithm such as total number of strategy flips, the mean number of generations before a strategy flip, and the mean magnitude of a strategy flip. All of these figures are for the mixed microhabitat usage trait.

**Fig 9.**
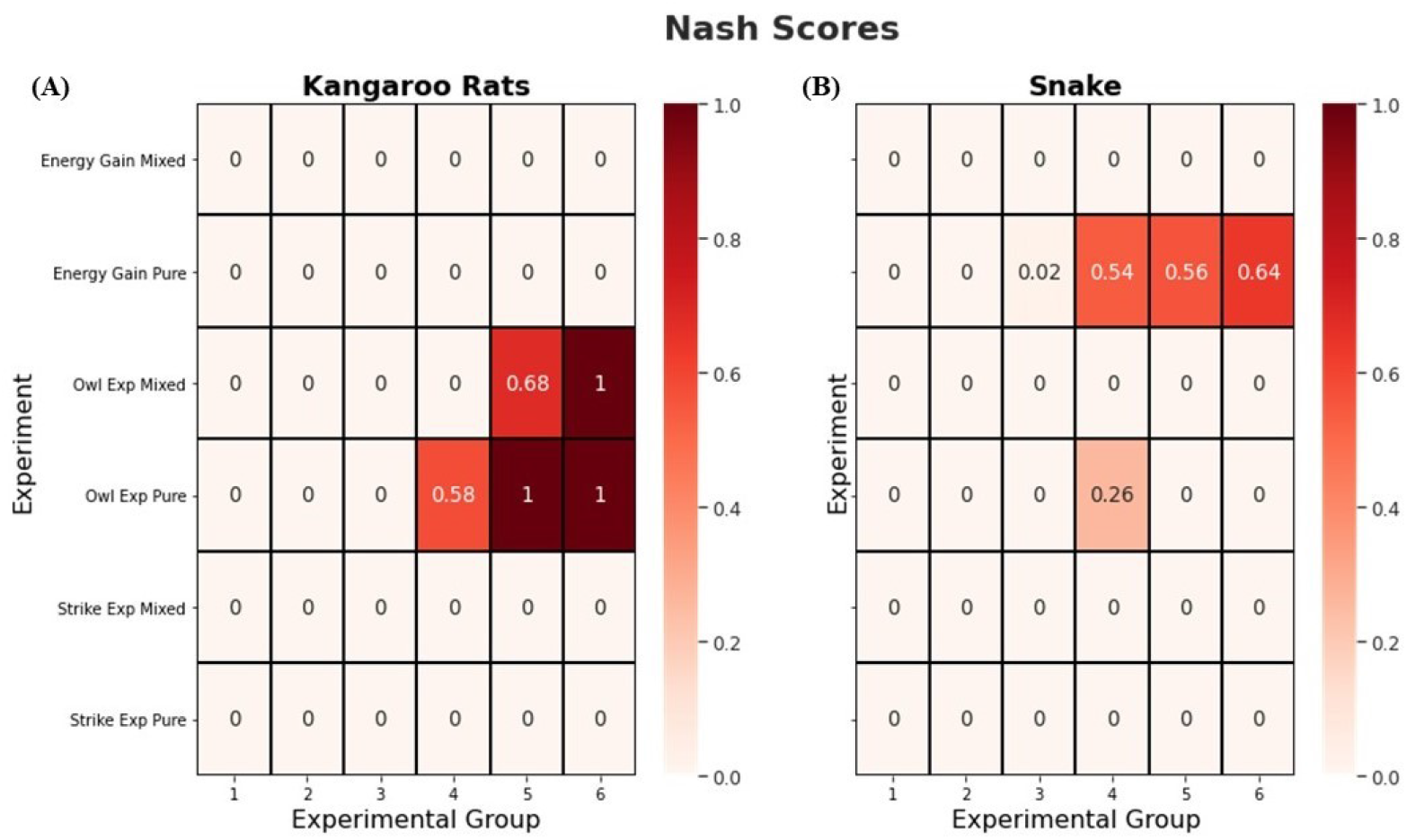
Results of the Nash Score analysis where each cell represents the Nash Score for that specific experimental group associated with the experimental design. The Nash Score is the ratio of simulations that reached equilibrium over the total number of simulations run for a given experimental design. A Nash Score of 1 means that 100% simulations completely reached equilibrium and the associated microhabitat preference strategy is an evolutionarily stable strategy and can’t be invaded by other strategies throughout evolutionary time. (A) shows Nash Score values for the kangaroo rat population and (B) shows the same information but for the rattlesnake population.

### 2.1 Case Study 1: Differing Foraging Energy Gains for Kangaroo Rats in Microhabitats

This set of simulations involved progressively increasing the amount of energy individual kangaroo rats gain from foraging in the open compared to the bush microhabitat (Table 2), while keeping the effects of predation constant. When modeling microhabitat preference as a pure trait, in lower ordered experimental groups where the benefit of foraging in a particular microhabitat is greater than the cost of foraging (experimental groups 1, 2 and 3. respectively) the kangaroo rat population is distributed evenly in both microhabitat types (Figure 5A), with no statistical difference between these three experimental groups (Figure 4A). After experimental group 4, the kangaroo rat population adopts a dominant preference of the open occurs (Figure 5A) and there is no statistical difference in preference between experimental groups 4,5, and 6 (Figure 4A). As the disparity of energy gain in one microhabitat versus the other grows large enough (experimental groups 4, 5, and 6 Table 2) and one of the microhabitats can no longer provide enough energy to sustain the metabolic costs (Table 1), then prey preference shift completely toward the habitat with greater resources (Figure 5A). However, before this tipping point (experimental groups 2,3 Table 2), the resource disparity has no effect kangaroo rat microhabitat preference (Figure 4A)

The rattlesnakes, however, showed a more gradual transition to preferring the open in experimental groups 1,2, and 3. In experimental group 4 the population completely adopts a microhabitat preference for the open (Figure 5B) and we see that experimental groups 4, 5, and 6 are statistically identical (Figure 4B). The rattlesnake population microhabitat preference for the pure experiment showed high levels of evolutionary stability, as interpreted by our ESI. Experimental group 4 and 5 have a Nash Score greater than 0.5 (Figure 9B) meaning that over half of the simulations run with these specific experimental parameters reached an equilibrium state. In contrast, the kangaroo rat populations in this set of simulation has a Nash Score of 0 (Figure 9A), with a large number of strategy flips and high variance for experimental groups 3-6 (Figure 7A). The kangaroo rat mean number of generations before a strategy flip and the mean magnitude of the strategy flip (Figure 7B, C) are relatively low in experimental groups 3,4,5, and 6 compared to experimental groups 1 and 2 further demonstrating that they are switching strategies often.

The results of simulations when modeling microhabitat preference as a mixed trait were generally similar, with both kangaroo rats and rattlesnakes adopting an increased preference for the open in experimental groups with a higher disparity in food resources, but maintain a limited preference for bush microhabitat persisted for all experimental groups (Figure 6A, B). Nearly all experimental groups are statistically different than one another for the kangaroo rat population except for experimental group 1 and 2 (Figure 4C). The Nash Score analysis shows that neither population is close to reaching an ESS in this case study as all experimental groups have a Nash Score of 0 (Figure 9A, B).

A discrepancy between pure versus mixed strategies can be observed in experimental groups 3-6 (Table 2), kangaroo rats retain more preference for the bush habitat than in the pure strategy simulations (Figure 5A, B, 6A, B). Additionally, mean number of generations before a strategy flip for the kangaroo rats (Figure 8B) is lower compared to the mean generations before a strategy flip of the other case studies (Figure 8E, H) and the mean magnitude of the strategy flip (Figure 8C) is low in these experimental groups. This indicates that the kangaroo rat population is not altering microhabitat preference over time as drastically, primarily using the open (Figure 6A), while occasionally using lower quality bush microhabitats. For the rattlesnakes, it is optimal to partake in some degree of prey behavioral management by changing behavioral strategies as seen in the slight preference toward the bush (Figure 6B).

### 2.2 Case Study 2: Differing Strike Success of Rattlesnakes in the Bush Microhabitats

To explore the impact of varying rattlesnake strike success, each subsequent experimental group gradually increased the probability of rattlesnakes successfully striking prey in bush microhabitats while decreasing strike success in the open (Table 3). When microhabitat preference is modeled as a pure trait, kangaroo rats gradually shifts preferences toward the open across experimental groups, with the mean difference between experimental group 1 to experimental group 6 is − 0.188 (Figure 4E). The magnitude of microhabitat preference shifts for the rattlesnake population by comparison is much smaller, as the largest difference between experimental groups is from group 1 to 5 with about a 0.05 preference shift for bush microhabitats. Neither the kangaroo rat population or the rattlesnake population microhabitat usage strategy are evolutionary stable as the Nash Score of all experimental groups is 0 (Figure 9). Both populations also have a high mean magnitude of strategy flips (Figure 7F) and the mean generations before a strategy flip is constant which indicates both populations are switching strategies often through evolutionary time and the rattlesnake strike success probability has little effect on driving microhabitat usage toward equilibrium.

This same set of conditions resulted in very different outcomes when microhabitat preference was modeled as a mixed trait. Rather than trending slightly toward a bush preference (Figure 5D), the mixed trait rattlesnake population tends to prefer the open (Figure 6D), with a mean difference of − 0.138 between experimental groups group 1 and 6 (Figure 4H). This increased preference toward the open appears to reach a limit as the mean difference between experimental group 5 and 6 is only − 0.008 and is less statistically significant compared to the differences of the other experimental groups (Figure 4H). In contrast, kangaroo rat preferences were similar when modeled as either mixed or pure traits, although they did show an increased propensity for the open at higher experimental groups (Figure 6C). For both populations, these strategies are not evolutionary stable (Figure 9). There is evidence that increasing the strike success probability does have a somewhat stabilizing effect on microhabitat preference, as the mean magnitude of strategy flips does decrease in subsequent experimental groups for both populations (Figure 8F). Experimental group 5 and 6 have an increased mean generations before a strategy flip (Figure 8E), meaning populations are not switching strategies as often compared to the control group (experimental group 1).

### 2.3 Case Study 3: Introducing Owls

Introducing owls into the system had a large and significant effect on the microhabitat usage patterns of rattlesnakes and kangaroo rats. For the pure microhabitat usage strategy, both kangaroo rats and rattlesnakes had similar responses to the introduction of owls (Figure 5E, F). Experimental group 1 and 2 show a gradual shift in microhabitat preference toward the bush with a mean microhabitat usage difference of 0.07 for the kangaroo rat population (Figure 4I) and 0.05 for the rattlesnakes (Figure 4J). Once the proportion of owls is equal to rattlesnakes (experimental group 4), an increased preference toward the bush is seen in both populations. Experimental group 3 to 4 has a mean difference of microhabitat preference of 0.327 for kangaroo rats (Figure 4 I) and 0.322 for rattlesnakes (Figure 4 J).

When modeling preference as a pure trait, kangaroo rats preference toward the bush the bush starts approaching equilibrium in experimental group 4 with a Nash Score of 0.58 and then becomes an ESS with a Nash Score of 1.0 for experimental groups 5 and 6 (Figure 9A). The rattlesnake population does not reach equilibrium in any of the experimental groups for this case study. Experimental group 4 has a Nash Score of 0.26, meaning that the population was starting to reach equilibrium, but the subsequent experimental groups 5 and 6 did not (Figure 9B). For the rattlesnake population, the total number of strategy flips (Figure 7G) dramatically drops from experimental group 3 to 4, but then increases again for experimental group 5 and 6. The mean generations before a strategy flip (Figure 7H) decreases from experimental group 4 to 5, indicating that there is more fluctuations in microhabitat usage strategies over time as the total number of rattlesnakes in the population decreases (Table 4).

Modeling microhabitat preference as a mixed strategy trait yields similar results to the pure owl experiment as when owls are the more abundant predator this causes a significant preference toward utilizing bush microhabitats; however, there are subtle differences in these two experiments. In the pure owl experiment, both populations adopted a 100% preference for the bush when owls are the more abundant predator (Figure 5E, F). In contrast, in the mixed experiment, when owls are dominant, kangaroo rats will prefer to use bush habitats approximately 90% of the time and rattlesnakes prefer the bush about 83% of the time. Experimental group 4, 5, and 6 are not significantly different in microhabitat preference (Figure 4L).

Another difference between the pure and mixed experiment is that in the mixed experiment, kangaroo rat microhabitat preference reaches an evolutionary stable equilibrium in later experimental groups. Experimental group 5 has a Nash Score of 0.68 and reaches complete equilibrium at experimental group 6 with a Nash Score of 1 (Figure 9A). The rattlesnake population again showed no evidence of approaching an evolutionary stable equilibrium. The total number of strategy changes for the rattlesnakes has a non-linear trend and there is actually an increase in strategy flips when comparing experimental group 4 and 5 (Figure 8G) and a slight increase in the mean magnitude of strategy flips from experimental group 4 to 5 and 5 to 6 (Figure 8I), indicating that rattlesnakes become more unpredictable in landscape usage at these higher experimental groups.

## 3 Discussion

### 3.1 The Nash Score and Expanding Agent-based Modeling Methodology

One of our primary modeling objectives was to integrate well known analysis techniques from evolutionary game theory into an agent-based framework to increase the standardization and interpretability of future simulation studies using agent-based models. Concepts such as behavioral strategies, pay off scores, pure or mixed strategies, and the ESS are all concepts developed for game theory that we integrated as inputs into our agent-based framework. Our model outputs summarize the optimal behavioral strategy in a given set of conditions, and formulated a new method for determining when the system reaches an ESS.

The Nash Score conceptually is analogous to the ESS as it describes when a behavioral trait transitions from cycling through many strategies over evolutionary time due to frequency dependent selection, to being completely stable from generation to generation. The dynamic of cyclical fluctuations of behavioral strategies over time has been observed in other agent-based predator prey models [21] and is due to adaptation of two trophic levels interacting and co-evolving with one another. This is also known as an evolutionary “chase” [1], as both predators and prey display bi-stable cyclical trends, with an evolutionary time lag in the predator population.

The notion of an ESS finds application in agent-based systems employing genetic algorithms are applied. When a population’s behavioral trait remains constant over the run time of a simulation, it signifies that any potential mutant strategies are suboptimal and unable to become fixed within the population. In mathematical terms, the ESS in an agent-based framework categorized by the Nash Score is quantified by a system transition from a bistable equilibrium to a unistable equilibrium for a given population. Because it is a continuous metric, the Nash Score also allows us to quantify if the system is approaching equilibrium and provides methods to assess equilibrium of populations independently of one another as certain selection pressures may impact one population’s behavioral strategies more than others.

However, one key difference between the Nash Score and traditional ESS analysis is the two populations (kangaroo rats and snakes) behavioral strategies are evaluated independently, thus one population may reach an ESS but the other may not. This is a key difference to keep in mind as our method more depicts whether the scenario will lead the population to an ESS versus in traditional game theory finding the ESS tells you the conditions of stability for both populations together. Thus the two concepts could have different application purposes and are not be mutually exclusive.

The model finds application in understanding frequency dependent behavioral strategies in particular, as the system will oscillate when the behavioral trait of one population is frequency dependent on the other populations choices, and the behavioral strategies of individuals in the given agent population. A Nash Score of *<* 1 depicts conditions fitness of the trait is going to be frequency dependent on the antagonist population (predator or prey). For the kangaroo rats, the fitness for microhabitat preference will be negatively frequency dependent as more kangaroo rats adopt a certain strategy, this will increase the fitness of the rattlesnakes to adopt the same microhabitat preference which then makes the rare trait have higher fitness, leading to arms race dynamics. This is inversely true for the rattlesnakes. If too many snakes adopt a certain microhabitat strategy, it will cause the other population to switch strategies. This is how trait oscillations form as an arms race between the two populations ensues.

However, when certain ecological conditions outweigh the fitness impacts of this arms race, certain microhabitat preference traits may stabilize which means that outside selection pressures outweigh the predator prey arms race. This showcases the utility of the Nash Score as it is capable of describing conditions where arms race dynamics dominate the system. When the Nash Score is 0, the frequency dependent predator prey arms race is the primary factor in the behavioral trait evolution; versus when other ecological conditions (such as owls) lead to trait stability and outweighs the arms race (Nash Score = 1) (Figure 9A).

This modeling framework gives researchers a new computational tool to describe the stability and trajectories of many simulations that would be difficult to analyze visually. For the case studies described, we ran a total of 1800 simulations and the Nash Score and the other metrics provided by the point classification algorithm provide a tool for understanding simulation trajectories without having to analyze each simulation individually which greatly reduces the analysis complexity. This makes it easier to assess the implications of how changes in ecological conditions impact overall community dynamics and behavioral trait evolution for future studies and formulate an objective narrative based on the results of many simulations.

### 3.2 Case Study 1: Differing Foraging Energy Gains for Kangaroo Rats in Microhabitats

One of the objectives of all of our experimental designs was to determine the ecological factors that most influence shifts and stability in the microhabitat preferences of a predator-prey system. For the first case study, increased energy gain heterogeneity, our findings add nuance to the general principle that predator microhabitat preferences track prey preferences, and prey balance predation risk with energy gain. In our analysis, we were able to identify a tipping point of the system between these two dynamics, suggesting environmental conditions shift the balance in power between predator and prey in terms of microhabitat preferences. A tipping point is when small changes in external environmental conditions cause large and abrupt changes to individuals’ behaviors, interactions among group members, and therefore how the group functions of the system [17]. Thus our analysis shows that there is a duality between these two hypotheses rather than a general rule of all systems.

Based off the pure strategy microhabitat usage, for kangaroo rats under the pure strategy, we identified there is a tipping point for the behavioral strategies. When there is a small gradient for heterogeneous foraging yields but both microhabitats provide a net energy gain, there is little to no behavioral changes for kangaroo rats at the population level, but we do observe behavioral changes in the rattlesnakes.

This can also be observed from the Nash Score analysis as after this tipping point is reached, the Nash Score begins to approach 1 for the rattlesnake population, although it never actually reaches the ESS. The Nash Score for the kangaroo population remains at 0 for all experimental groups. What this indicates from an ecological perspective is the kangaroo rats are becoming predictable in their microhabitat usage as one of the microhabitats no longer covers the cost of foraging, leading to stability in predator microhabitat usage strategies as the monopolize using the open rather than the bush. This may seem counter intuitive because the direct selection pressure of the tipping point is on the kangaroo rats, however, this leads to behavioral trait stability to the predators rather than the prey.

The previous finding along with simulations modeling microhabitat preference as a mixed trait underscore the importance of prey to be able to use microhabitats that may provide temporary spatial refuge from predators even at the cost of less or no energy gain. The rattlesnakes have a Nash Score of zero across all experimental groups which indicates they are switching behavioral strategies due to the kangaroo rats being less predictable in their microhabitat usage strategies. Overall, what this case study indicates is there are environmental conditions where prey predominately drive the behavioral strategies of the system as the predator tracks their landscape usage, versus conditions when prey are predictable and the predator monopolizing part of the landscape drives behavioral microhabitat usage patterns.

### 3.3 Case Study 2: Differing Strike Success of Rattlesnakes in Bush Microhabitats

From the Nash Score analysis, increased strike success heterogeneity only led to further arms race dynamics as there is no ESS and the Nash Score values are zero. Increasing the probability of a rattlesnake strike success in bush microhabitats had little effect on increasing preference toward the bush of the rattlesnake population in the pure experiment. There is even a direct conflicting effect in the mixed experiment as rattlesnakes have to adopt an open preference and have to switch microhabitat usage strategies often through evolutionary time despite being at a disadvantage in this microhabitat. The data of our experiments shows rattlesnakes must prefer the open microhabitat at high rates despite strike performance benefits from bush microhabitats because their increased killing efficiency in the bush drives kangaroo rats to the open. Thus, increased specializing in performance benefits for one microhabitat over another without some outside ecological pressure such as owls in the open or an increased foraging gain for kangaroo rats in the bush, leads the specialization to actually be maladaptive.

### 3.4 Case Study 3: Introducing Owls

The results from this case study indicate that the introduction of a predator specializing in foraging in open microhabitats (owls) will cause a shift in preference toward using the bush. Our results also imply that there can be a microhabitat preference tipping point when the number of owls exceeds that of rattlesnakes. If the trait is a pure strategy, then preference for bush microhabitats becomes an ESS for the kangaroo rats. The snakes adopt the same strategy, but it is not an ESS for snakes, suggesting that it is occasionally optimal for snakes to forage in the open, thus increasing selection pressure to drive kangaroo rats toward the bush.

One of the main goals of our study was to confirm the validity of our model by comparing it to a previous game theory model on kangaroo rat and rattlesnake predator prey interactions [5]. Our findings were similar to the traditional game theory model, which found that it is an ESS for both kangaroo rats and rattlesnakes preferring the bush when the probability of being captured by an owl in the open outweighs the probability of being capture by a snake in the bush. However, the rattlesnake population in our model never reaches equilibrium which could be because of low population sizes in conditions that should be evolutionary stable as the population briefly starts approaching equilibrium in experimental group 3 (Figure 9B) with a Nash Score of 0.26 but subsequent experimental groups have a Nash Score of 0. When the population size is low, even one member of the population having a different microhabitat preference will have a larger effect on the populations mean microhabitat preference due to a low sample size, thus making equilibrium harder to reach.

When microhabitat preference is modeled as a mixed trait, the owl simulations suggest there is a limit for the rattlesnake population’s bush preference regardless of the number of owls present. These results critique the idea that rattlesnakes and owls work synergistically [10] and supports the idea that it is optimal for rattlesnakes to distribute selection pressure in the open a minority of the time to monopolize prey in the bush.

This supports the idea that predators work to distribute “fear” across a landscape [7, 13, 14], however, owls will shift the theoretical fear distribution. It is evolutionary advantageous for rattlesnakes to switch behavioral strategies after a certain number of generations (Figure 8H) to increase the probability of a strike from either a rattlesnake or an owl against a kangaroo rat in the open, showcasing the benefits of being a generalist predator compared to a specialist. When this probability is larger than the probability of a strike in the bush, this drives the kangaroo rat population primarily using the bush to be an evolutionary stable strategy. This means that owls primarily benefit rattlesnakes rather than vice versa as kangaroo rats reach an ESS for preferring bush microhabitats due to two predators in the open versus only having one type of predator in the bush.

The assumptions are model makes that may complicate this finding is that we assume owls are incapable of catching kangaroo rats under bushes which may not be ecologically true as some bush microhabitats may not provide the perfect canopy protection [10]. Other assumptions of our model that may change its findings are that interspecific competition behaviors between owls and rattlesnake have been found where some owls may predate rattlesnakes which would have an impact on their microhabitat preference [19]. One aspect of our experimental design that may shift results is that in experimental groups 5 and 6 the rattlesnake population is low which lessens statistical impact of our findings.

Across all of our case studies, our study supports the hypothesis that, without outside ecological pressures sustaining certain microhabitat preferences, predators will exhibit foraging behaviors that diffuse the probability of predation across a landscape [7] and predators and prey will display chase dynamics over evolutionary time [1]. Predator microhabitat preferences diffuse risk across the landscape even when predators are directly disadvantage in a microhabitat, as seen in the strike success experiment. Although these predator-prey dynamics have been recovered in other models and empirical studies, our modeling approach provides a nuanced understanding of how these dynamics are dependent on ecological circumstances and can identify tipping points in which the behavior of a population drastically shifts. Overall, the overarching conclusion from the case study our analysis suggests is that there are environmental conditions where predator prey systems are driven by prey decisions, and conditions where predators drive system trajectories. This point can be seen as analogous to other complex behavioral system relationships such as employer-employee relations in economics where the scale of negotiating power based on the economic conditions and supply and demand [2].

## 4 Conclusion and Future Work

The most practical extension of this project is to leverage these methods as a way of building a bridge between hypothesis developed during field study collections, further integrating empirical data to estimate parameters, and testing hypothesis formed throughout an ecological field study trying to understand cryptic behaviors of organisms to formulate case studies akin to the ones demonstrated in this paper.

Along with our case studies, our modeling approach also allowed us to develop metrics that illustrate how these preferences shift through evolutionary time via metrics we developed such as the mean generations before a strategy flip, mean magnitude of the strategy flip, and the Nash score. Our model can also be applied to other biological systems, and other types of traits in order to develop a deeper understanding of predator-prey dynamics.

Future work in reference to the modeling framework would be to further develop several complementary extensions of our modeling approach. For example, our modeling framework would allow for randomly setting parameters in order to apply a regression-based approach to understanding how these ecological forces interact with one another and impact microhabitat preference. Some of the case studies showed non-linear trends in microhabitat preference shifts across the experimental groups due to trade offs between predation risks and foraging gain. Some of these limits may be shifted or overcome when introducing multiple ecological factors. For example, in our strike success study, rattlesnakes show only a modest increase in preferences for bush microhabitat, despite their greatly increased strike success. Their preference could be greatly increased if we also introduced owls into the landscape, or increased foraging gain potential in the bush for kangaroo rats.

Another potential ecologically relevant extension would be to include within-generation behavioral learning of predator and prey in order to model the impact of behavioral experience in these populations. This has been successfully accomplished in other models such as [28] and would better incorporate how ontogeny effects landscape usage over evolutionary time. Applying genetic algorithms or a learning algorithm to the owl population as well would perhaps provide a more accurate model of predator interference and would address the potentially limiting assumption that owls cannot hunt in bush microhabitats.

The benefit of taking an agent-based modeling approach is that it is relatively easy to incorporate more complexity. The design and analysis of our initial case studies provide a standardized framework to obtain several important insights on how microhabitat preferences of predator and prey population shift as a result of ecological complexity. Our approach allows us to measure the magnitude trait shifts and their evolutionary stability. These tools in conjunction with empirical studies can help better understand how ecology influences the evolution of certain traits and can be applied to most other biological systems.

## Supplementary Material

### Sensitivity Analysis

We conduct a “One-factor-at-a-time” (OAFT) protocol for sensitivity analysis of our model [18]. The base parameter being manipulated can be found in Table 1 and the values they are changed to can be found in the legend of Figure 10 next to the parameter name. Ten simulations are run and the mean microhabitat preference is calculated for both kangaroo rats and rattlesnakes. From this analysis, we find that overall, the system is the most sensitive to changes in landscape parameter “open to bush microhabitat proportion” compared to other parameters, especially in the mixed model (Figure 10 B). This makes sense as preference would likely be correlated with the abundance of a given microhabitat. Both the mixed and the pure model are relatively robust to parameters involving game theoretic components such as energy cost and energy gain, population sizes, and overall landscape size in the x and y direction (Figure 10).

**Fig 10.**
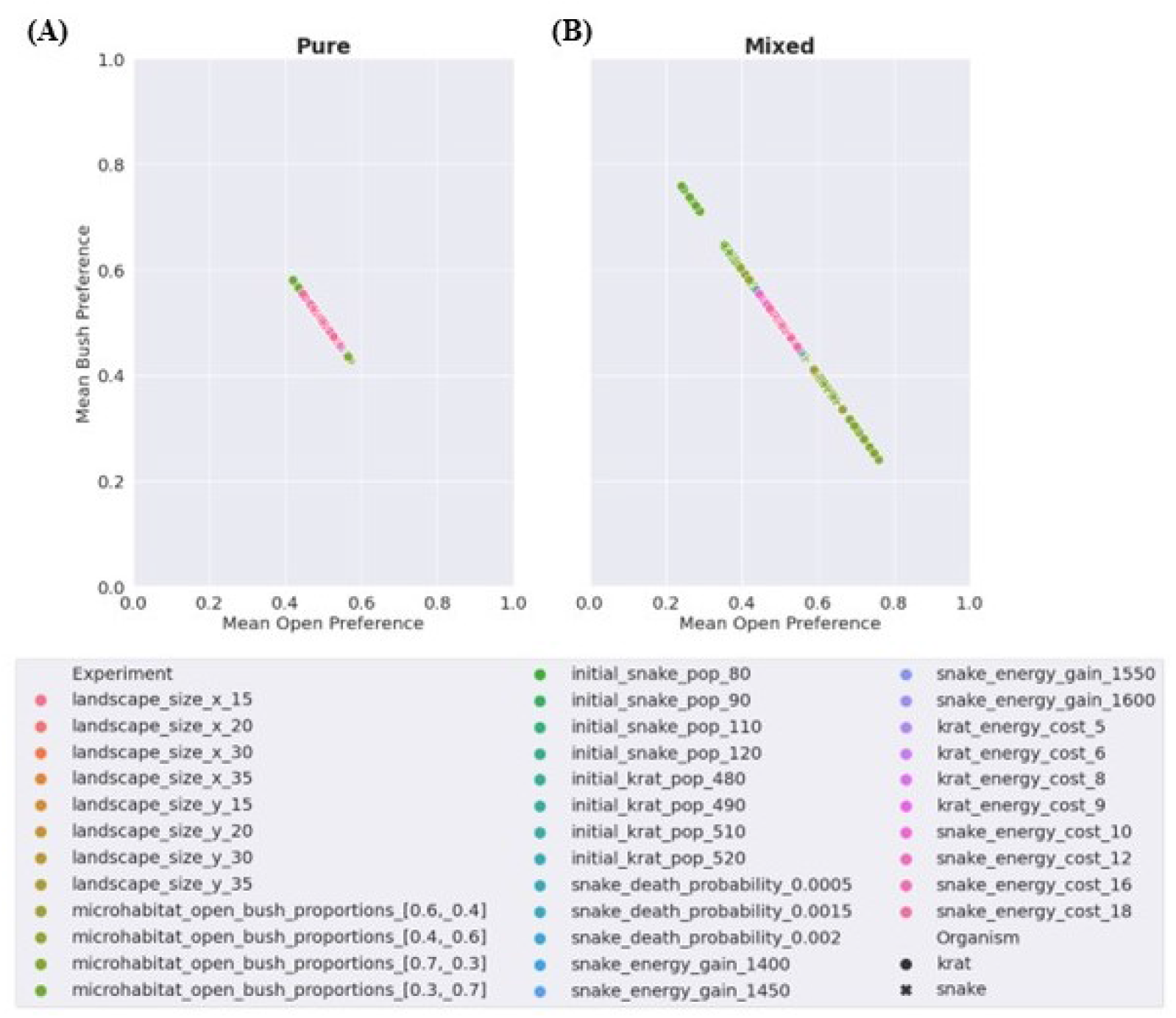
Results of sensitivity analysis of the microhabitat preferences of kangaroo rats and rattlesnakes. Each point is the mean microhabitat preference of ten simulations for the population with the given parameter being changed. Color represents the parameter being changed and the value it is being changed to. Circles indicate results for kangaroo rats (krats) while x’s represent rattlesnake results. See Table 1 for the base parameters that are being manipulated.

### Optimal Point Classification Accuracy

To obtain metrics such as number of strategy flips, mean generations before a strategy flip, mean magnitude of a strategy flip, and Nash Score we developed an algorithm developed to classify points as local maxima or local minima in the per generation time series from single simulations. To evaluate the performance of this algorithm, I manually classified local optimality points from a subset of two different time series, and then applied the classification algorithm to form a confusion matrix and calculate metrics such as accuracy, precision, and recall.

The first was the snake population generations 1500− 1665 from simulation 0 of pure owl experimental group 2 (Table 4) which had 63 optimal points. The algorithm had an accuracy of 0.95, a precision of 0.92, and a recall of 0.94. The second subset was from the kangaroo rat population of the mixed owl experimental group 1 (Table 4) from generations 9500 − 9998 which had 19 optimal points. Our algorithm had an accuracy of 0.95, a precision of 0.17, and a recall of 0.05. This showcases that the algorithm in some time series has propensity for false negatives in certain time series regions. Overall, our algorithms performance can range greatly and further work may be to make the algorithm less conservative as it can miss optima points in some time series. Despite this, we still see clear trends in our metrics provided by this algorithm as seen in Figure 7 and Figure 8 likely due to the sheer amount of simulations run and the large number of generations for each simulation which could be statistically compensating for regions where the algorithm does poorly.

## Acknowledgments

A special thanks to Dr. Margaret Field Prof. Emerita, American Indian Studies at San Diego State for recommending the name Uumarrty for our software. Uumarrty is a Kumeyaay word which means “the gambling game”. We picked this name because the gambling aspect reflects the risk and reward trade off of organisms foraging in a game theory framework. We also wanted to pay homage to the native people of San Diego, the Kumeyaay tribe.

## Notes

### Competing Interest Statement

The authors have declared no competing interest.

### Summary of Updates

Updated title, revised typos, improved writing flow.

